# Redefining Ovarian Fibrosis Through Comparative Analysis of Collagen Architecture

**DOI:** 10.1101/2025.11.12.688021

**Authors:** Caroline J. Aitken, Lanna Kadhim, Alyssa Murray, David A. Landry

## Abstract

Ovarian fibrosis is a known pathology of reproductive aging, becoming a growing concern for infertility and complex ovarian diseases. In research, mouse and human ovary samples are utilized, though distinct differences between species warrant validation of architectural phenotypes to accurately define its pathology. Using polarized light microscopy and orientation analysis of collagen fibers in mouse and human ovaries, we define ovarian fibrosis as the accumulation and/or anisotropic organization of fibrillar collagen within the ovarian stroma and/or cortex.

## Introduction

Fibrosis is the excessive buildup of collagenous extracellular matrix (ECM), which can progressively cause significant scar tissue and loss of organ function[1]. The ovaries are particularly susceptible to fibrosis due to aging and subsequent changes in the inflammatory environment[2], causing shifts in immune cell profiles and myofibroblast activation[3]. Since ovarian fibrosis can entail damage to ovarian tissue structure and function, its implications in infertility and other ovarian diseases such as PCOS, endometriosis, and premature ovarian insufficiency have been a growing concern[4]. Additionally, there is compelling evidence that ovarian fibrosis contributes to ovarian cancer permissiveness and colonization, providing an optimal metastatic niche[5]. Due to the lack of mechanistic studies on ovarian fibrosis, current research is utilizing mouse models and human ovary samples to understand its pathology at the cellular and transcriptomic level.

Mouse models are valuable for studying fibrosis as they recapitulate the dynamic three-dimensional structures of the ECM and cell-cell communications involved in fibrotic processes[6]. Mouse models of ovarian fibrosis have been successfully developed using aging models of both immune deficient and immune compromised mice[3]. However, given the distinct structural and cyclical differences between mouse and human gonads, comparisons between histological phenotypes in mouse and human fibrosis require further investigation. Since fibrosis research is largely reliant on in vivo mouse models, understanding the differences between animal and human ovarian fibrosis is crucial for adequate interpretation of research findings and translational relevance[6]. This report aims to delineate species-specific phenotypes of ovarian fibrosis and highlight differences that will assist further research in this field.

## Results and Discussion

The architectural features of the fibrillar collagen matrix, specifically its density, thickness, and fiber characteristics (orientation and coherency) were assessed using Picrosirius Red (PSR) staining under polarized light microscopy (POL-PSR). This staining technique has previously been validated as a robust representation of the fibrillar collagen matrix within ovarian fibrotic tissue through comparative analyses with second harmonic generation (SHG) microscopy[3]. POL-PSR thus represents an effective, reliable, and cost-efficient alternative to SHG for the assessment of collagen fiber characteristics, while simultaneously providing additional information on fibrillar collagen thickness[7].

A marked qualitative difference in collagen staining intensity was observed between species, which was further explored through analysis of birefringence patterns (Figure 1). The birefringence originates from the molecular organization of collagen fibers, whose ordered triple helices refract polarized light differently by orientation; the parallel alignment of PSR molecules along these fibers amplifies this intrinsic property through their linear dichroism[8]. This property enables the visualization of fiber thickness and organization: thick, tightly packed fibers appear orange to red, whereas thin, loosely organized fibers appear green to yellow[9]. Historically, the birefringence colours produced by PSR-POL have been interpreted as indicators of collagen type, with type I collagen displaying orange-red hues and type III collagen appearing green-yellow[8]. However, this interpretation has been challenged by recent evidence showing a weak correlation between birefringence colour and collagen type in rat tendons[7], indicating that PSR-POL colour should primarily be considered a reflection of fiber thickness rather than collagen type.

**Figure 1:**
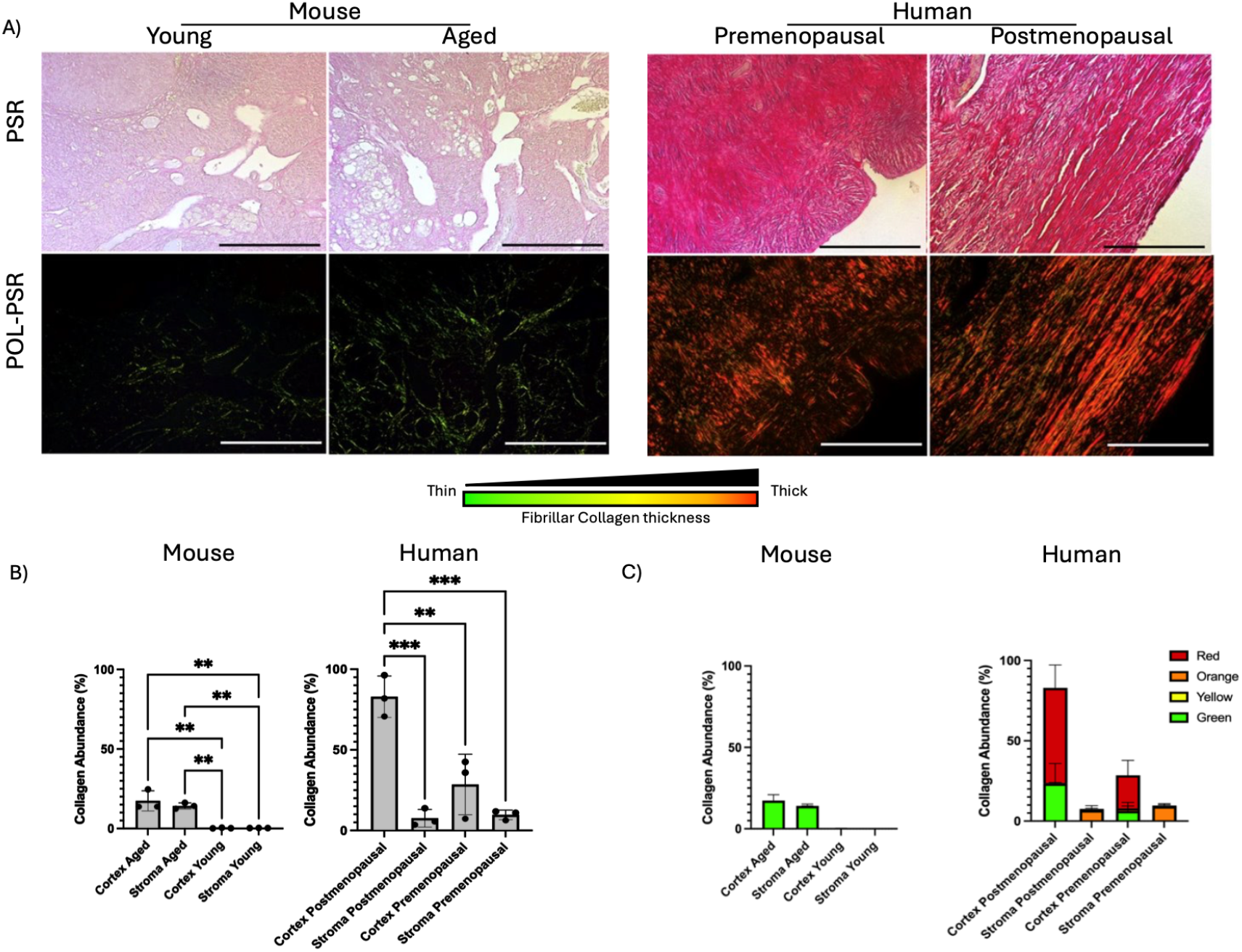
Mouse and human ovarian collagen abundance: **A**. Collagen deposition from PSR-stained ovaries of young (6 months) and aged (18 months) mice compared to PSR-stained premenopausal and post-menopausal human ovaries, visualized under brightfield in the top row, and polarized light (POL-PSR) in the bottom row. Collagen thickness is determined by HUE, where thin fibers are represented from green to red pixels. **B**. Quantification of collagen abundance by total HUE pixels in the ovarian cortex and stroma of mice and humans. Total positive pixels were normalized to the total pixel count and averaged within each group. **C**. Quantification of collagen thickness by HUR pixel frequency and saturation in ovarian cortex and stroma of mice and humans (n=3). Data are means ± SEM (\**P <* 0.05, \*\**P <* 0.01, and \*\*\**P <* 0.001). Scale bars 200µm.

In this study, we analyzed PSR staining of mouse and human ovaries under polarized light microscopy to quantify birefringence and fibrillar collagen organization in non-fibrotic and fibrotic ovaries at two defined depths: the cortex (50 µm below the ovarian surface epithelium, OSE) and the stroma (1 mm below the OSE) (Figure 1A). This approach allowed us to compare collagen distribution and organization patterns between species and across aging. Our quantitative analysis of birefringence images revealed an age-associated increase in fibrillar collagen deposition in both mouse and human ovaries (Figure 1B), predominantly composed of thin fibers in mice and thick fibers in humans (Figure 1C). In aged mice, collagen accumulation was evident in both the cortex and stroma. In contrast, in human ovaries, collagen accumulation was restricted to the cortical region, with no detectable difference between premenopausal and postmenopausal stroma. Thus, collagen deposition in aging human ovaries appears spatially confined to the cortex, unlike the more diffuse pattern observed in mice. These findings are consistent with Landry et al., who reported age-related increases in thin collagen fibers throughout the mouse ovary[3]. To our knowledge, collagen fiber thickness in aging human ovaries has not been previously quantified; however, McCloskey et al. described a pronounced anisotropic organization of fibrillar collagen fibers in the ovarian cortex following menopause, as shown by SHG microscopy[10], and Vaishnav et al. showed a similar phenotype using a digital platform for analysis of collagen structure[11].

We observed similar organizational differences in human ovaries; however, the anisotropic organization was limited to the cortex region of post-menopausal women (Figure 2). This species-specific distinction in collagen organization is illustrated in Figure 2A, which presents representative images of collagen fiber architecture. The directional distribution of fibers within each image was quantified using orientation histograms (Figure 2B). In these histograms, an isotropic fiber network appears as a flat distribution, whereas a network exhibiting preferential alignment displays a distinct peak in the dominant direction. Collectively, these analyses demonstrate that age-associated ovarian fibrosis in mice does not lead to the same level of anisotropic organization seen in postmenopausal human ovaries. This observation is supported by collagen coherency measurements derived from the orientation analysis (Figure 2C & 2E), which indicate high fiber alignment in postmenopausal human ovaries but low coherency in aged mouse ovaries. When coherency was compared between stroma and cortex by age, no significant differences were observed in mouse ovaries (Figure 2D), regardless of region or age. In contrast, human ovaries showed significantly higher cortical coherency compared to the stroma in postmenopausal women (Figure 2F). Interestingly, while the stroma remained unchanged across menopausal status, a strong trend (P = 0.052) suggested lower anisotropic organization in the ovarian cortex of premenopausal compared to postmenopausal women.

**Figure 2:**
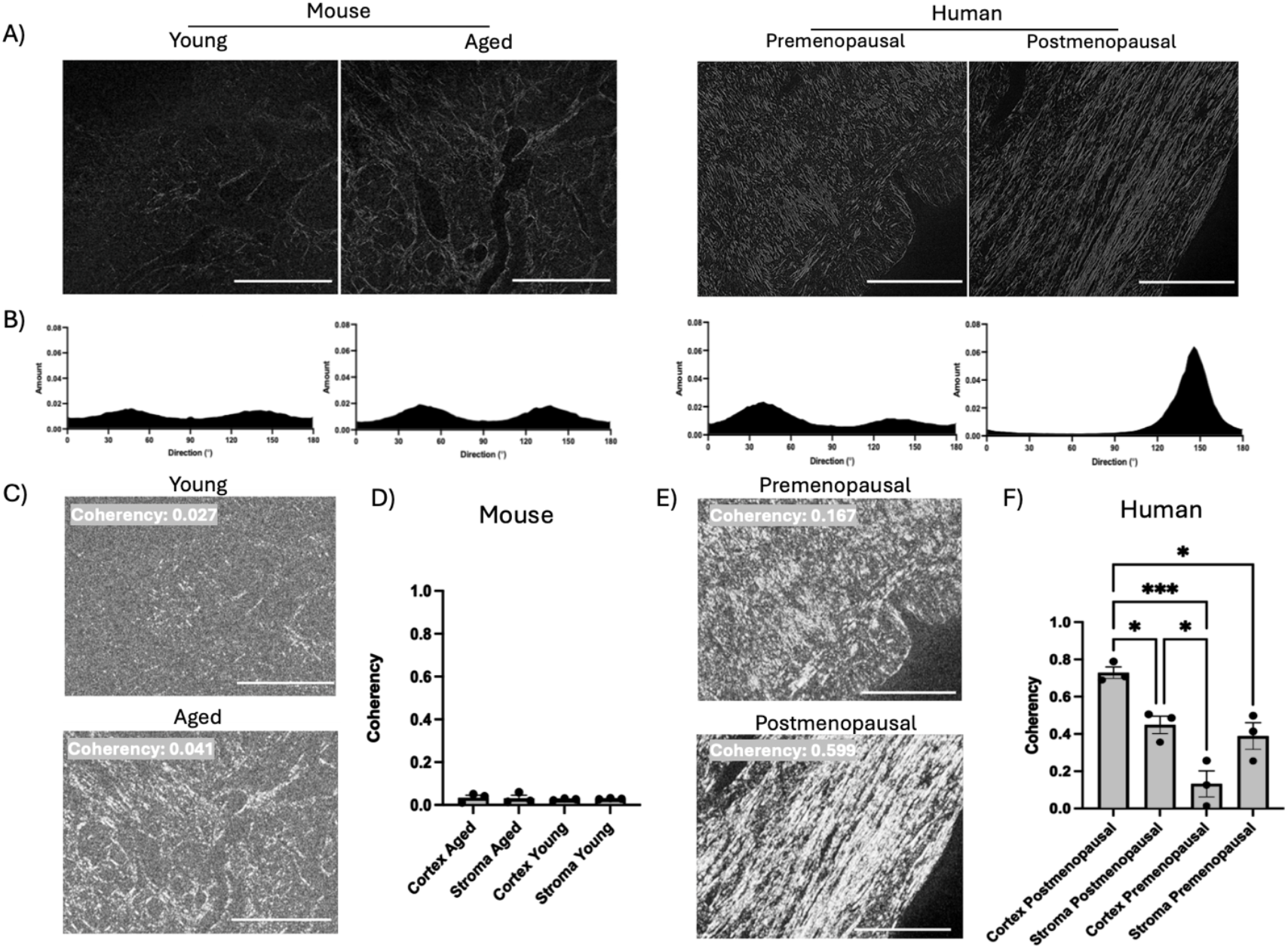
Mouse and human ovarian collagen orientation and coherence: **A**. Collagen organization under polarized light from PSR-stained young (6 months) and aged (18 months) mouse ovaries, and human pre- and post-menopausal ovaries. **B**. Representative orientation histograms showing the distribution of collagen fiber angles (pronounced peaks denote dominant alignment directions, while flatter distribution indicates isotropic organization). Fiber coherency visualized and quantified by measuring the saturation of local gradients in mouse (**C**) and human (**E**) ovaries. High saturation (coherency values near 1.0) indicates parallel fiber alignment, whereas low saturation (values near 0.0) indicates perpendicular or disorganized fibers. Bar graphs represent the comparison of coherency measurements within ovarian cortex and stroma of mice (**D**) and humans (**F**) (n=3). Data are means ± SEM (\**P* < 0.05, \*\**P* < 0.01, and \*\*\**P* < 0.001). Scale bar 200µm.

Taken together, the observed differences between aged mouse and human ovaries may likely reflect underlying variations in collagen subtype composition and fibrillar organization. These differences in collagen subtype regulation have been previously reported, as COL3A1 gene networks differ between mouse and human, supporting the possibility of species-specific pathways in development and resolution of fibrosis[12]. On the contrary, some mouse models of fibrosis have been shown to have similarities to some molecular features of human fibrotic diseases, with variability depending on the organ, etiology, and genetic background[13], [14]. Consequently, while murine models provide valuable mechanistic insight, their limitations must be recognized when extrapolating to human ovarian fibrosis, particularly regarding collagen subtype dynamics and matrix organization during aging.

## Conclusion

Physiological factors such as ovulation pattern and frequency, along with differences in cortical cellularity, immune cell distribution, and the duration of reproductive aging, may further contribute to species-specific patterns of ovarian remodeling. In humans, less frequent ovulation, prolonged exposure to postmenopausal hormonal changes, and a higher baseline collagen content may together predispose the cortex to anisotropic organization and deposition with aging. In contrast, in mice, repeated ovulatory injury combined with faster tissue repair and lower stromal density may drive a distinct fibrotic trajectory characterized by more diffuse collagen deposition. These distinctions suggest that both the mechanisms underlying fibrosis and the kinetics of its resolution may differ between species. Future studies integrating comparative proteomic and extracellular matrix analyses of mouse and human ovaries will be critical to identify the specific collagen subtypes, structural ECM components, and molecular pathways involved, thereby clarifying the origin and evolution of age-associated ovarian fibrosis. Such comparative approaches will also be essential for interpreting findings from experimental mouse models and for assessing potential therapeutic intervention strategies. As a final remark, the definition of ovarian fibrosis should be refined to encompass not only the accumulation of collagen but also the alterations in its spatial organization, particularly the anisotropic arrangement observed in the human ovarian cortex. Based on our findings, we propose the following definition: *‘‘Ovarian fibrosis is defined as the age- or injury-associated accumulation and/or anisotropic organization of fibrillar collagen within the ovarian stroma and/or cortex, leading to altered extracellular matrix architecture*.*’’*

## Materials and Methods

This study was performed under a protocol approved by the Animal Care Committee of the University of Ottawa and conducted in accordance with the guidelines of the Canadian Council on Animal Care. Mice were housed under controlled environmental conditions with 12-hr alternating light/dark cycles, with free access to water and food. Young (6 months old) and aged (18 months old) female C57BL/6J mice were purchased from the Jackson Laboratory. Mice were euthanized based on age, and ovaries were collected for histological analysis. Human ovarian tissues from 3 premenopausal and 3 postmenopausal women were obtained in accordance with the Ottawa Health Science Network Research Ethics Board protocol 20180168-01H and generously donated by Dr. Barbara Vanderhyden [11].

Histopathologic assessment of murine and human ovaries was performed using 5 μm sections of formalin-fixed paraffin embedded tissue. Briefly, tissues were incubated in 10% (w/v) neutered formaldehyde buffer (NFB) for 24h at room temperature and transferred to 70% ethanol to be paraffin-embedded at the Louise Pelletier Histology Core Facility at the University of Ottawa. Sections were prepared at a thickness of 5 μm and stained with PSR. Slides were imaged at 20x under bright-field (PSR) and polarized light (POL-PSR) using a Zeiss AXIO Imager M2 with a linear polarizer and an Axiocam 105 color camera (Zeiss, Oberkochen, Germany; CBIA core Facility at the University of Ottawa).

For fibrillar collagen quantification, polarized images were analyzed using FIJI (ImageJ). For each image, we performed the same subtraction operation to remove any background and non-collagen birefringence. We then determined the hue (color) of each pixel within the subtracted image. To determine the value of each hue pixel, the hue component was retained, and a histogram of hue frequency and saturation was obtained from the resolved 8-bit images (which contain 256 colors). We used the following hue definitions as previously described[3]; red 2-9 and 230 – 256, orange 10-38, yellow 39-51, and green 52-128. The hue between 129-229 consists of non-birefringent tissue and is labeled as no collagen. The number of pixels within each hue range was determined and expressed as a percentage of the total image area (200 um x 200 um) in pixels. To measure the orientation and coherence, images were analyzed using the OrientationJ plugin to calculate the directional coherency coefficient. A coherency coefficient close to 1 indicates a strongly coherent orientation of the local fibers in the direction of the long axis. A coherency coefficient close to zero denotes no preferential orientation of the fibers[15].

All statistical analyses were performed using Prism 10 software (GraphPad Software Inc., La Jolla, CA, USA). Datasets were compared using analysis of variance with Tukey’s multiple comparisons posttest. All data are presented as means ± SEM.

## Data and Materials Availability

All data needed to evaluate the conclusion in the manuscript are present in the manuscript.

## Author Contribution

Conceptualization: DAL; Acquisition of data: CJA, LK, AM, DAL; Interpretation: CJA, AM, DAL; Writing original draft: CJA, AM, DAL; Review & editing: CJA, LK, AM, DAL; Supervision and final approval: DAL

## Acknowledgment

We are grateful to Dr. Barbara Vanderhyden for providing the human ovarian PSR-stained slides. We would also like to acknowledge the Cell Biology and Image Acquisition Core (RRID: SCR_021845) funded by the University of Ottawa and the Canada Foundation for Innovation, the Animal Care and Veterinary Services Core Facility at the University of Ottawa, and the Louise Pelletier Histology Core Facility at the University of Ottawa.

## References

[1] N. C. Henderson, F. Rieder, and T. A. Wynn, “Fibrosis: from mechanisms to medicines,” Nature, vol. 587, no. 7835, pp. 555–566, Nov. 2020, doi: 10.1038/s41586-020-2938-9.

[2] J. Wu, Y. Liu, Y. Song, L. Wang, J. Ai, and K. Li, “Aging conundrum: A perspective for ovarian aging,” Front. Endocrinol., vol. 13, p. 952471, Aug. 2022, doi: 10.3389/fendo.2022.952471.

[3] D. A. Landry, E. Yakubovich, D. P. Cook, S. Fasih, J. Upham, and B. C. Vanderhyden, “Metformin prevents age-associated ovarian fibrosis by modulating the immune landscape in female mice,” Sci. Adv., vol. 8, no. 35, p. eabq1475, Sept. 2022, doi: 10.1126/sciadv.abq1475.

[4] D. A. Landry, H. T. Vaishnav, and B. C. Vanderhyden, “The significance of ovarian fibrosis,” Oncotarget, vol. 11, no. 47, pp. 4366–4370, Nov. 2020, doi: 10.18632/oncotarget.27822.

[5] H. Fujimoto et al., “Tumor-associated fibrosis: a unique mechanism promoting ovarian cancer metastasis and peritoneal dissemination,” Cancer Metastasis Rev., vol. 43, no. 3, pp. 1037–1053, 2024, doi: 10.1007/s10555-024-10169-8.

[6] J. Padmanabhan, Z. N. Maan, S. H. Kwon, R. Kosaraju, C. A. Bonham, and G. C. Gurtner, “In Vivo Models for the Study of Fibrosis,” Adv. Wound Care, vol. 8, no. 12, pp. 645–654, Dec. 2019, doi: 10.1089/wound.2018.0909.

[7] C.M. López De Padilla, M. J. Coenen, A. Tovar, R. E. De la Vega, C. H. Evans, and S. A. Müller, “Picrosirius Red Staining: Revisiting Its Application to the Qualitative and Quantitative Assessment of Collagen Type I and Type III in Tendon,” J. Histochem. Cytochem., vol. 69, no. 10, pp. 633–643, Oct. 2021, doi: 10.1369/00221554211046777.

[8] L. C. Junqueira, G. Bignolas, and R. R. Brentani, “Picrosirius staining plus polarization microscopy, a specific method for collagen detection in tissue sections,” Histochem. J., vol. 11, no. 4, pp. 447–455, July 1979, doi: 10.1007/BF01002772.

[9] D. Dayan, Y. Hiss, A. Hirshberg, J. J. Bubis, and M. Wolman, “Are the polarization colors of picrosirius red-stained collagen determined only by the diameter of the fibers?,” Histochemistry, vol. 93, no. 1, pp. 27–29, 1989, doi: 10.1007/BF00266843.

[10] C. W. McCloskey et al., “Metformin Abrogates Age-Associated Ovarian Fibrosis,” Clin. Cancer Res. Off. J. Am. Assoc. Cancer Res., vol. 26, no. 3, pp. 632–642, Feb. 2020, doi: 10.1158/1078-0432.CCR-19-0603.

[11] H. T. Vaishnav et al., “BRCA mutation alters the stromal landscape in normal ovaries,” Oct. 03, 2025, bioRxiv. doi: 10.1101/2025.10.01.679597.

[12] L. Wang et al., “Differences between Mice and Humans in Regulation and the Molecular Network of Collagen, Type III, Alpha-1 at the Gene Expression Level: Obstacles that Translational Research Must Overcome,” Int. J. Mol. Sci., vol. 16, no. 7, pp. 15031–15056, July 2015, doi: 10.3390/ijms160715031.

[13] A.K. Martínez et al., “Mouse Models of Liver Fibrosis Mimic Human Liver Fibrosis of Different Etiologies,” Curr. Pathobiol. Rep., vol. 2, no. 4, pp. 143–153, Dec. 2014, doi: 10.1007/s40139-014-0050-2.

[14] L. Walkin et al., “The role of mouse strain differences in the susceptibility to fibrosis: a systematic review,” Fibrogenesis Tissue Repair, vol. 6, no. 1, p. 18, Sept. 2013, doi: 10.1186/1755-1536-6-18.

[15] E. Fonck et al., “Effect of aging on elastin functionality in human cerebral arteries,” Stroke, vol. 40, no. 7, pp. 2552–2556, July 2009, doi: 10.1161/STROKEAHA.108.528091.

